# Aerodigestive sampling reveals altered microbial exchange between lung, oropharyngeal, and gastric microbiomes in children with impaired swallow function

**DOI:** 10.1101/476580

**Authors:** Claire Duvallet, Kara Larson, Scott Snapper, Sonia Iosim, Ann Lee, Katherine Freer, Kara May, Eric Alm, Rachel Rosen

**Affiliations:** Department of Biological Engineering, MIT, Cambridge, Massachusetts; Center for Microbiome Informatics and Therapeutics, MIT, Cambridge, Massachusetts; Aerodigestive Center, Division of Gastroenterology, Hepatology and Nutrition, Boston Children’s Hospital, Boston, Massachusetts; Division of Gastroenterology, Hepatology and Nutrition, Boston Children’s Hospital, Boston, Massachusetts; Division of Pulmonary Medicine, Boston Children’s Hospital, Boston, Massachusetts

**Keywords:** Aerodigestive microbiomes, respiratory, oral, and gastric microbiomes, gastroesophageal reflux, impedance, aspiration, video fluoroscopic swallow study

## Abstract

1

**Background:** Children with oropharyngeal dysphagia have impaired airway protection mechanisms and are at higher risk for pneumonia and other pulmonary complications. Aspiration of gastric contents is often implicated as a cause for these pulmonary complications, despite being supported by little evidence. The goal of this study is to determine the relative contribution of oropharyngeal and gastric microbial communities to perturbations in the lung microbiome of children with and without oropharyngeal dysphagia and aspiration.

**Methods:** We conducted a prospective cohort study of 220 patients consecutively recruited from a tertiary aerodigestive center undergoing simultaneous esophagogastroduodenoscopy and flexible bronchoscopy. Bronchoalveolar lavage, gastric and oropharyngeal samples were collected from all recruited patients and 16S sequencing was performed. A subset of 104 patients also underwent video fluoroscopic swallow studies to assess swallow function and were categorized as aspiration/no aspiration. To ensure the validity of the results, we compared the microbiome of these aerodigestive patients to the microbiome of pediatric patients recruited to a longitudinal cohort study of children with suspected GERD; patients recruited to this study had oropharyngeal, gastric and/or stool samples available. The relationships between microbial communities across the aerodigestive tract were described by analyzing within- and between-patient beta diversities and identifying taxa which are exchanged between aerodigestive sites within patients. These relationships were then compared in patients with and without aspiration to evaluate the effect of aspiration on the aerodigestive microbiome.

**Results:** Within all patients, lung, oropharyngeal and gastric microbiomes overlap. The degree of similarity is the lowest between the oropharynx and lungs (median Jensen-Shannon distance (JSD) = 0.90), and as high between the stomach and lungs as between the oropharynx and stomach (median JSD = 0.56 for both; p = 0.6). Unlike the oropharyngeal microbiome, lung and gastric communities are highly variable across people and driven primarily by person rather than body site. In patients with aspiration, the lung microbiome more closely resembles oropharyngeal rather than gastric communities and there is greater prevalence of microbial exchange between the lung and oropharynx than between gastric and lung sites (p = 0.04 and 4×10^−5^, respectively).

**Conclusions:** The gastric and lung microbiomes display significant overlap in patients with intact airway protective mechanisms while the lung and oropharynx remain distinct. In patients with impaired swallow function and aspiration, the lung microbiome shifts towards oropharyngeal rather than gastric communities. This finding may explain why antireflux surgeries fail to show benefit in pediatric pulmonary outcomes.

## 2 Introduction

The economic and social impact of oropharyngeal dysfunction and aspiration is well known in the adult stroke population; adults with oropharyngeal dysfunction are at greater risk of pneumonia than those without [1]. Little is known about aspiration-related lung disease in children, though recent studies suggest that up to 10% of all pneumonia hospitalizations in pediatrics are related to aspiration [2]. Clinicians often assume these pneumonias result from the aspiration of refluxed gastric contents and frequently treat these children with antireflux surgery, fundoplication [3, 4]. Despite this common surgical practice [5, 6], there are no pediatric studies which conclusively show improved pulmonary outcomes after fundoplication, suggesting that the respiratory symptoms seen in aspirating patients may not be related to aspiration of gastric contents [7, 8, 9, 10, 11]. An alternative hypothesis is that aspiration-related respiratory symptoms may result from aspirated oropharyngeal contents. To test this hypothesis, we determined the microbial signatures of the lungs, stomach, and oropharynx in children with and without oropharyngeal dysphagia (i.e. with and without impaired airway protective mechanisms) to determine the relative contributions of the oropharyngeal and gastric microbiomes to the lung microbiome. We quantified the relationships between communities both within and across patients by calculating the beta diversity between samples and by defining individual OTUs exchanging between sites in multiple patients.

Previous studies have shown that the mouth, upper respiratory tract, and lung microbiota contain similar microbes, and that upstream oral communities seed downstream sites (e.g. lungs and stomach) [12, 13, 14]. However, there is little consensus on whether there exists a distinct or “core” lung microbiome that is consistent across people [13, 15, 16, 17]. Most studies, however, agree that the lung microbial communities share taxa with the oral microbiome, but that there are some bacteria present in lung communities whose abundances cannot be traced solely to the mouth [12, 13, 16, 18].

While the importance of oropharyngeal flora in seeding the lungs has been heavily studied in ICU settings [19, 20, 21], the role of oropharyngeal-lung flora exchange in otherwise healthy children with isolated swallowing dysfunction is unknown. Furthermore, studies investigating the relationships between microbial communities across the aerodigestive tract have not examined how microbes exchange between the stomach and lungs, and how this exchange relates to clinical factors such as aspiration and gastroesophageal reflux.

If the lung microbiome of aspirating patients exhibits more exchange with the oropharynx than the stomach, this could provide evidence for why anti-reflux surgery is not helpful in patients with aspiration-related respiratory symptoms. Furthermore, a shift in the lung microbial communities toward an oropharyngeal population could not only result in overt pneumonia but may also have more subtle, pro-inflammatory effects [22]. Finally, if there is a unique aerodigestive microbial signature in aspirating patients, microbial profiling may be helpful as a diagnostic tool for oropharyngeal dysphagia or in follow-up validation cohorts to identify subsets of patients who may be at higher risk for pneumonia.

## 3 Methods

### 3.1 Patient cohort and sample collection

We conducted a prospective cross sectional cohort study of children ages 1-18 undergoing bronchoscopy and esophagogastroduodenoscopy (EGD) for the evaluation of chronic cough. Patients with gastrostomy or nasogastric tubes, a history of gastrointestinal surgery, or antibiotic use at the time of sample acquisition were excluded. The study was approved by the Boston Children’s Hospital Institutional Review Board and informed consent was obtained from all patients/parents. Information about the patient demographics and symptoms are included in Table 1 and Supplementary Table 1.

**Table 1:**
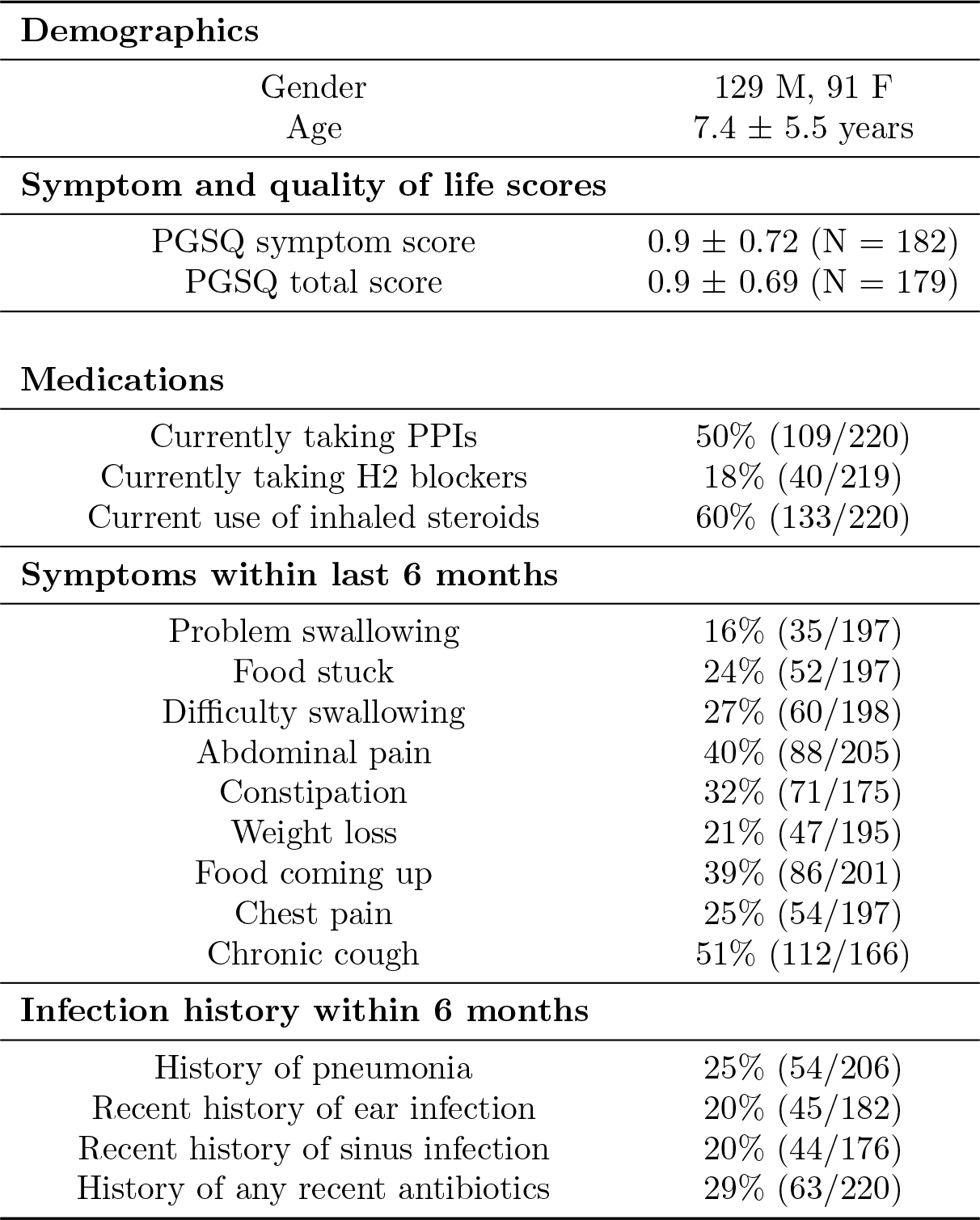
Patient demographics. While all patients were given questionnaires, not all patients completed the answers to all questions.

We first performed brushing of the posterior tongue to obtain oropharyngeal samples, placing the brush in TE buffer at −80C. Second, the bronchoscopy and bronchoalveolar lavage (BAL) was performed through an endotracheal tube in distal airways of the right middle lung or the most visually inflamed lung. Finally, gastric sampling was performed during the EGD. The endoscope was advanced, without suctioning, immediately into the stomach where the gastric fluid was suctioned into a sterile leukitrap. A minimum of 1 cc of gastric and lung fluid were collected and transferred to −80C.

All patients undergoing bronchoscopy had a triad of samples collected: oropharynx, gastric fluid, and BAL (Table 2 and Supplementary Table 2) [14]. To contextualize our findings, we also compared the aerodigestive microbiome of pediatric patients with suspected GERD who had oropharyngeal, gastric and/or stool microbiome samples collected. Additionally, many of the BAL samples were unable to be sequenced due to low DNA content. Thus, not all 220 patients have sequencing data for the same combination of samples. Tables with additional information describing the samples collected from each patient and which samples were used in each analysis are available at https://github.com/cduvallet/aspiration-analysis-public/final/supp_files.

**Table 2:** Number of patients with each combination of body sites sequenced.

|  | Number of patients |
| --- | --- |
| BAL, gastric fluid, and oropharyngeal swab | 66 |
| BAL and gastric fluid | 22 |
| BAL and oropharyngeal swab | 7 |
| Gastric fluid and oropharyngeal swab | 45 |
| Oropharyngeal swab and stool | 20 |
| BAL only | 6 |
| Gastric fluid only | 12 |
| Oropharyngeal swab only | 37 |
| Stool only | 5 |
| Total patients | 222 |

### 3.2 Multichannel intraluminal impedance with pH (pH-MII)

A subset of patients had pH-MII testing at the discretion of the patient’s primary gastroenterologist. Acid reflux episodes were defined as episodes detected by the impedance (MII) sensors with associated drop in pH to < 4; non-acid episodes did not have the associated drop. The percentage of time that reflux was in the proximal/distal esophagus was calculated by dividing the sum of the bolus clearance times in the proximal/distal esophagus by the total study duration. The percentage of full column reflux events was defined as the percentage of the total reflux events that reached the proximal two impedance sensors (i.e., the proximal most impedance channel) [23].

### 3.3 Oropharyngeal dysphagia assessment

A subset of the patients included in this study had a videofluoroscopic swallow study (VFSS) to asses swallow function and were divided into two groups (normal swallow function and aspiration/penetration). Because patients with penetration on VFSS have similar pulmonary symptoms and respond similarly to thickening as patients that aspirate, we included patients with aspiration and penetration in one group.

### 3.4 Sample processing and sequencing

Oropharyngeal swabs, BAL, and gastric fluid samples suspended in Tris-Saline buffer were centrifuged for 3 minutes at 10,000 rcf prior to DNA isolation. DNA was extracted from the sample pellet with the Qiagen DNeasy PowerSoil Kit as described by the manufacturer, with the following modifications: protein precipitation in one step using 100 *μ*L of each C2 and C3 solutions, and column centrifugation at 10,000 rcf for 10 minutes. Library preparation and sequencing was performed in two batches at the Broad Institute. 515F and 806R primers were used to amplify a ∼250bp region from the V4 region of the microbial 16S gene. Paired-end sequencing was performed on a MiSeq (175bp paired). Patients with multiple samples had all of their respective samples sequenced in the same batch.

### 3.5 Microbiome data processing and community analyses

Paired end reads were merged using USEARCH `-fastq_mergepairs` and truncated to 200 bp. Reads with more than 2 expected errors were discarded. Operational taxonomic units (OTUs) were clustered at 99% similarity and assigned taxonomy using the RDP classifier (c = 0.5) [24]. All quality filtering and OTU calling steps were performed with an in-house pipeline (https://github.com/thomasgurry/amplicon_sequencing_pipeline).

Beta diversity was calculated with an in-house implementation of the Jensen-Shannon distance (JSD) [25], which is calculated by taking the square root of the Jensen-Shannon divergence. The Jensen-Shannon divergence is a measure of divergence between distributions accounting for both presence and abundances of organisms and which deals well with the compositionality of microbiome data; the square root of the Jensen-Shannon divergence converts this into a distance metric, which are the values we report here [25, 26]. JSD values close to 1 indicate that two communities are very different, while values close to 0 correspond to more similar communities. Although this metric has been used broadly in microbiome research [27, 28], we also include results with an alternative beta diversity metric, the Bray-Curtis distance, in the Supplementary Figures. Only samples which were sequenced in the same batch were considered in cross-patient comparisons. Differences in overall community structure across sites was assessed using the PERMANOVA test as implemented in scikit-bio v 0.4.2 (`skbio.stats.distance.permanova`).

Alpha diversities were calculated on the raw OTU counts using Python’s `alph.shannon`, `alph.chao1`, and `alph.simpson` functions in `skbio.diversity.alpha`. Differential abundance analysis between aspirators and non-aspirators was performed on the relative abundances of OTUs and genera using a Kruskal-Wallis test implemented in Python’s `scipy.stats.mstats` module (function `kruskalwallis`, a non-parametric test and an implementation which accounts for ties [29]). P-values were corrected for multiple hypothesis testing with the `multipletests` function from `statsmodels.sandbox.stats.multicomp`, with the Benjamini/Hochberg correction (`method = ‘fdr_bh’`). Corrections were performed separately for each aerodigestive site and taxonomic level.

### 3.6 Exchanged OTUs definition

To define exchanged OTUs, we used data from patients with all three sites sequenced (N = 66). For each OTU, we calculated the Spearman partial correlation 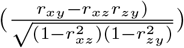 between its non-zero abundances in two sites, partialled on the third site (Scipy v 0.19.0 `stats.spearmanr`, [29]). P-values for each OTU were calculated as the percentage of null correlations larger than the observed correlation after shuffling abundances 2000 times. Only OTUs present in two sites in at least 10 patients were considered. OTUs with FDR-corrected q-value < 0.1 were defined as “exchanged” (`sandbox.stats.multicomp.multipletests with method=‘fdr_bh’`). To determine the statistical significance of the number of exchanged OTUs, we shuffled the patient IDs for each OTU in each site and re-defined “null” exchanged OTUs as described above.

### 3.7 Random Forest classifiers

We used Random Forest classifiers (scikit-learn v 0.18.1 `ensemble.RandomForestClassifier` with n_estimators=1000) for all supervised machine learning analyses [30]. For the classifier used to distinguish between aerodigestive sites (Figure 3B), we used 5-fold cross validation, ensuring that both samples from the same patient were in the same train or test split. For all other classifiers used to predict aspiration status, we performed a leave-one-out analysis. For each sample, we trained a model on all the other samples and used that model to predict the left-out sample’s label and label probability. Areas under the ROC curve (AUCs) and Fisher-pvalues were calculated based on these leave-one-out predictions using the `roc_curv, auce`, and `confusion_matrix` functions from Python’s `sklearn.metrics` module [30].

### 3.8 Availability of data and materials

Code to reproduce the analyses presented here are available at www.github.com/cduvallet/aspiration-analysis-public. The 16S sequencing data used in this study are available in the SRA repository at accession number SRP141148 and clinical metadata are available upon request from the corresponding author.

## 4 Results

Two hundred and twenty patients were included in the analysis (Tables 1 and 2; Supplementary Tables 1 and 2). The mean age of the patients was 7.4 ± 5.5 years. One hundred and nine out of 220 patients were taking proton pump inhibitors at the time of sampling. One hundred and four patients had a videoflouroscopic swallow study of which 47 (45%) had evidence of aspiration or penetration and 57 (55%) had normal swallow function. Of the 47 patients with aspiration or penetration, 26 patients had aspiration and 21 patients had isolated penetration. Of the patients with aspiration, 50% (n=13) aspirated thin liquids alone, 26.9% (n=7) aspirated thin and nectar consistency, 15.4% (n=4) aspirated thin, nectar and honey consistency and 7.7% (n=2) aspirated all textures including purees Twenty eight patients had pH-MII testing for gastroesophageal reflux at the time of sample collection. No relevant symptoms or clinical outcomes were significantly associated with aspiration status (Supplementary Table 1).

### 4.1 Aerodigestive microbiome across people

At the genus level, pediatric aerodigestive communities share many predominant members, including *Streptococcus*, *Prevotella*, *Haemophilus*, *Veillonella*, and *Neisseria* (Figure 1). However, despite genus-level similarities, OTU-level aerodigestive communities are distinct and highly variable across people. The overall community composition was significantly different between sites (PERMANOVA on JSDs between BAL, gastric fluid, and oropharyngeal samples in the two sequencing batches separately, p < 0.001, Figure 2A). Furthermore, lung communities were very different across people (median lung-lung JSD = 0.87) while oropharyngeal communities tended to be more similar (median oropharyngeal-oropharyngeal JSD = 0.59, Figure 2B).

**Figure 1:**
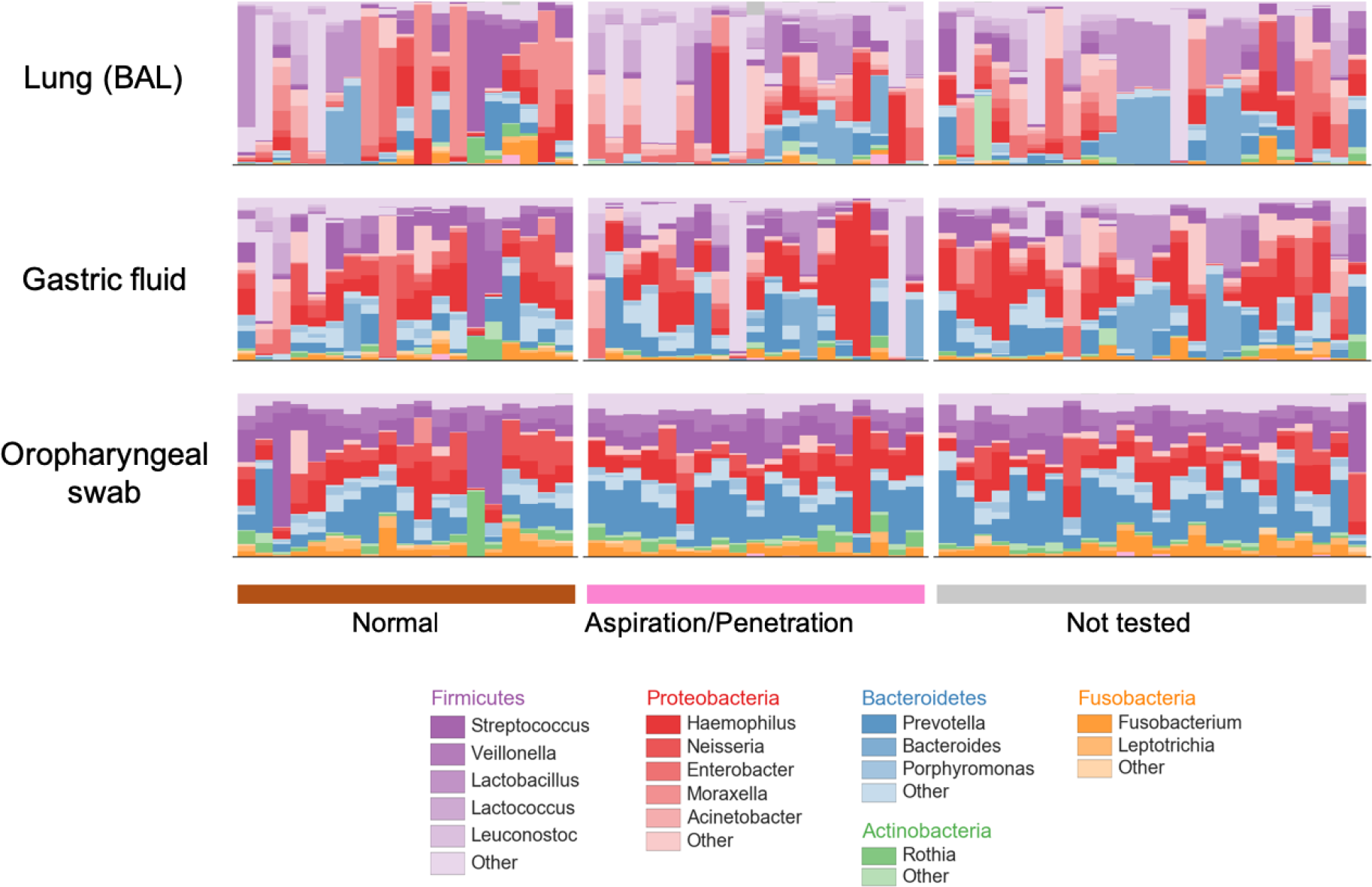
Aerodigestive communities have similar predominant genera. Bar plots showing relative abundances of aerodigestive microbiomes collapsed to the genus level for the 66 patients with all sequencing data from all three aerodigestive sites. Each column corresponds to one patient who had all three aerodigestive sites sequenced (N = 19 non-aspirators, 23 aspirators, 24 untested). Phyla in legend are those with mean abundance > 0.01 across all patients. Any other phyla are colored gray.

**Figure 2:**
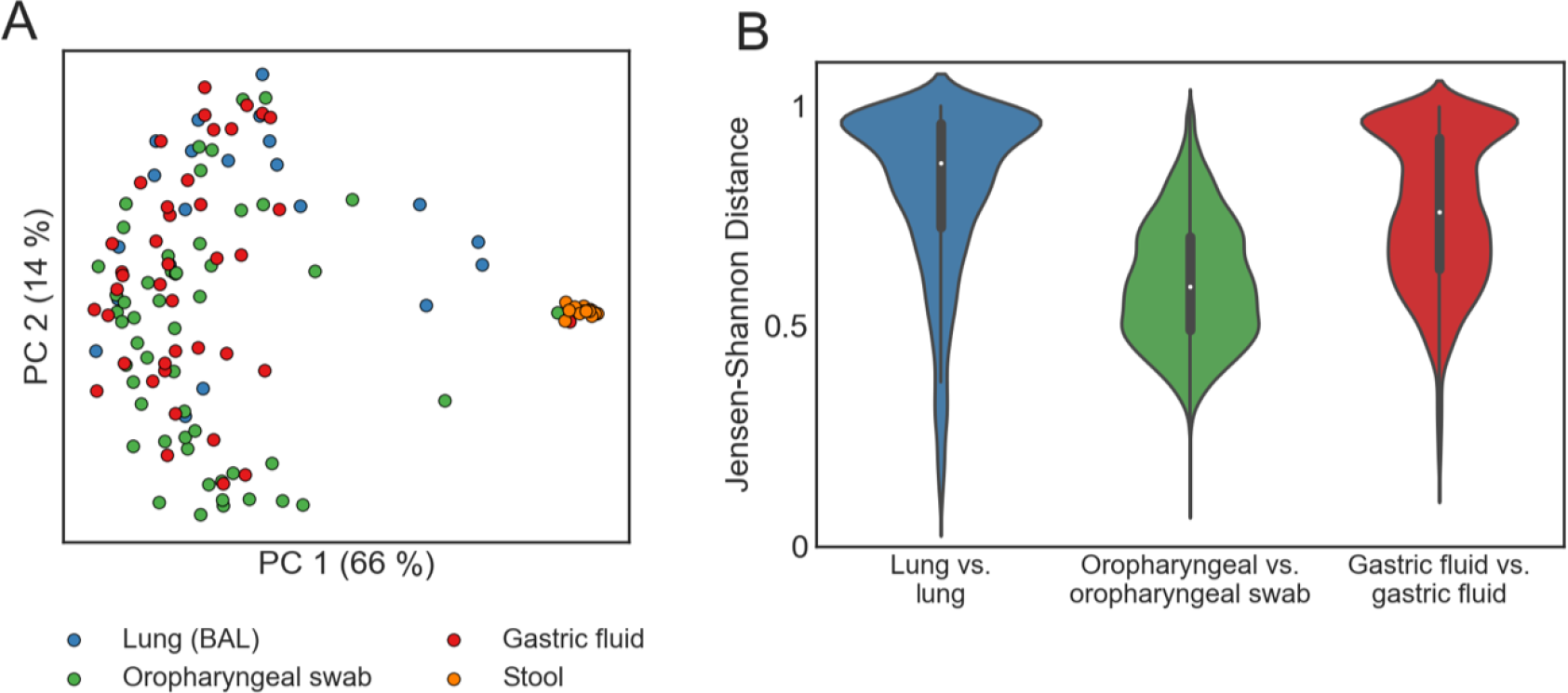
Lung and gastric communities are more variable across people than oropharyngeal communities. (A) PCoA plot of aerodigestive and stool microbial communities for all patients in the one sequencing batch (N = 21 BAL, 52 oropharyngeal swab, 43 gastric fluid, and 14 stool samples). The PCoA plot of the samples in the other sequencing batch are included in Supplementary Figure 1. (B) Violin plots of the Jensen-Shannon distance (JSD) between samples from the same site across different patients. A JSD close to 1 indicates that communities are very different (less similar). Supplementary Figures 4 and 5 show these results with the Bray Curtis distance metric instead of JSD.

**Figure 3:**
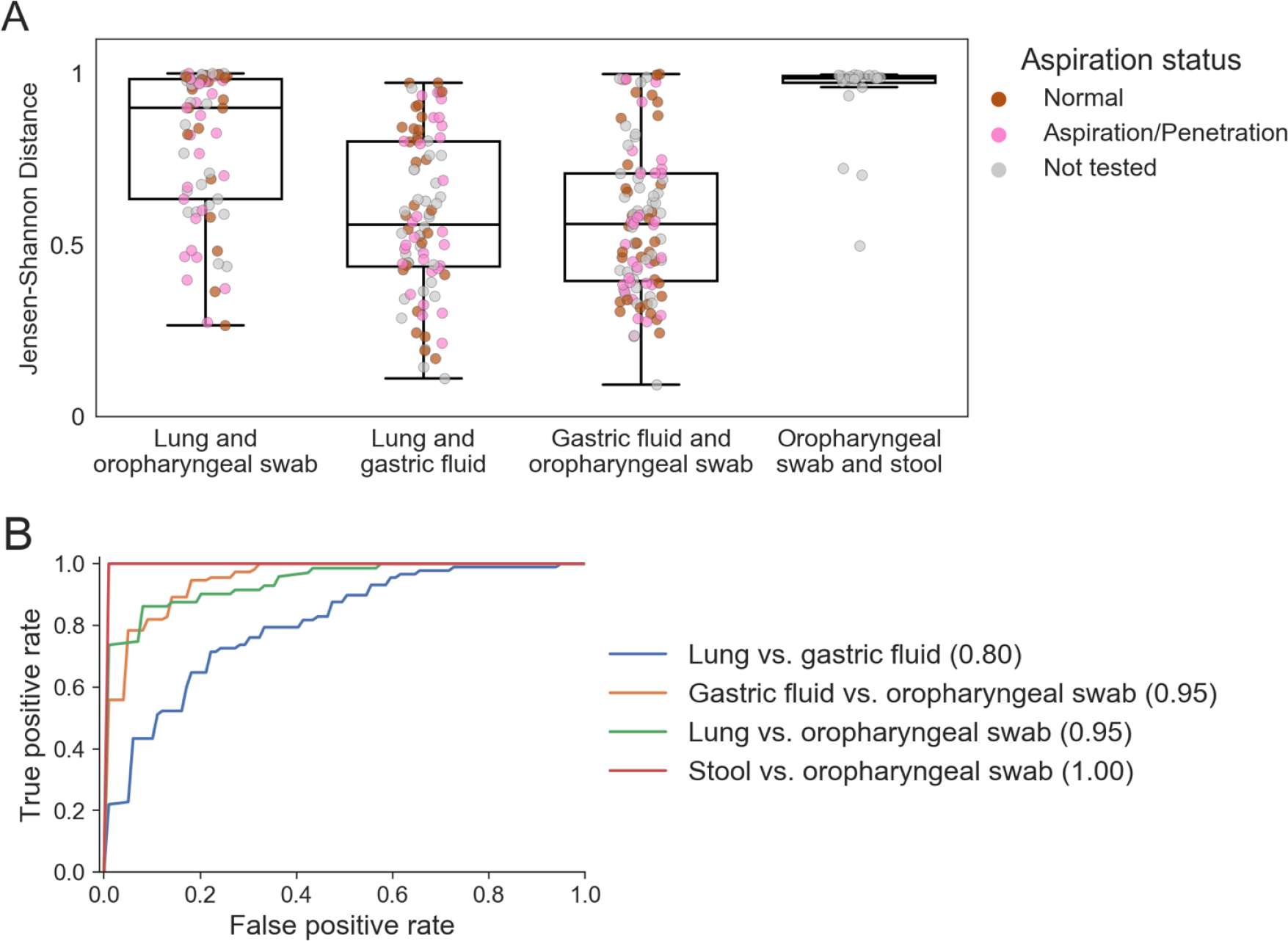
Within patients, aerodigestive communities are similar but lung and oropharynx remain most distinct. (A) Jensen-Shannon distances between samples from different sites from the same patient. Comparisons between stool and oropharynx are included to contextualize these results, as these are expected to be very different. All comparisons are significant (Wilcoxon rank sums test calculated with Python’s `scipy.stats.ranksums` function) except the lung and gastric fluid vs. gastric fluid and oropharyngeal swab beta diversities (p = 0.6). Lung and oropharyngeal vs. oropharyngeal and stool, p = 0.005. All other comparisons: *p* < 1 × 10^−8^. (B) ROC curve of classifiers distinguishing different aerodigestive sites. Mean areas under the ROC curve (AUCs) are reported in parentheses in the legend.

### 4.2 Aerodigestive microbiome within people

We compared aerodigestive communities within patients who had multiple sites sequenced (Table 2, Figure 3). Oropharyngeal and gastric fluid communities are similar within patients (median JSD = 0.56), reflecting that the mouth seeds the gastric microbiome [12, 13]. The majority of patients had very different lung and oropharyngeal communities (median JSD = 0.90), and these differences were significantly higher than either the lung-gastric fluid or gastric fluid-oropharyngeal beta diversities (p < 1 × 10^−8^, Figure 3A). Surprisingly, lung and stomach communities were as similar to each other as stomach and oropharyngeal communities (median JSD = 0.56 for both comparisons, p = 0.6).

We next identified specific microbes which exchange between aerodigestive sites within people. To do this, we reasoned that an actively exchanging microbe’s abundances in two sites should be correlated across patients (Supplementary Figure 2 and Methods). In other words, if an OTU is exchanged between two sites, if we observe that its abundance is low in both sites of one patient and high abundance in one site of another patient, then we would expect that its abundance in the second site of that second patient will also be high. We identified 13 OTUs exchanged between lung and oropharyngeal, 76 between gastric fluid and lung, and 117 between oropharyngeal and gastric fluid communities. These results were statistically significant: we found a maximum of 2 exchanged OTUs between sites in our null analysis. The low number of directly exchanged OTUs between the oropharynx and lungs supports the finding that these sites are more distinct than others in the aerodigestive tract. The lungs and stomach exchange fewer OTUs than the oropharynx and stomach even though they have comparable intra-patient similarities, suggesting that factors other than specific bacterial exchange contributes to the similarity between lungs and stomachs within patients.

Random Forest classifiers trained to distinguish between sites (ensuring that samples from the same patient were in the same train/test set) were able to identify a generalizable oropharyngeal microbial signature that distinguishes the oropharynx from other sites across people (AUC = 0.95 for both gastric fluid and lung comparisons, Figure 3B). Interestingly, when we compared within-patient similarities across sites to across-patient similarities for the same sites, we found that lung and stomach communities within patients were more similar than lungs across patients and than stomachs across patients (Figure 4, p < 1 × 10^−7^, Supplementary Table 3). Thus, while there exists a “core” oropharyngeal microbiome across people, lung and gastric communities are more variable and driven primarily by the person rather than body site. These results challenge the prevailing hypothesis that human-associated microbial communities are primarily driven by body habitat and instead suggest that patient-specific relationships may be equally, if not more, important in determining community structure in the aerodigestive microbiome [31, 32, 33].

**Figure 4:**
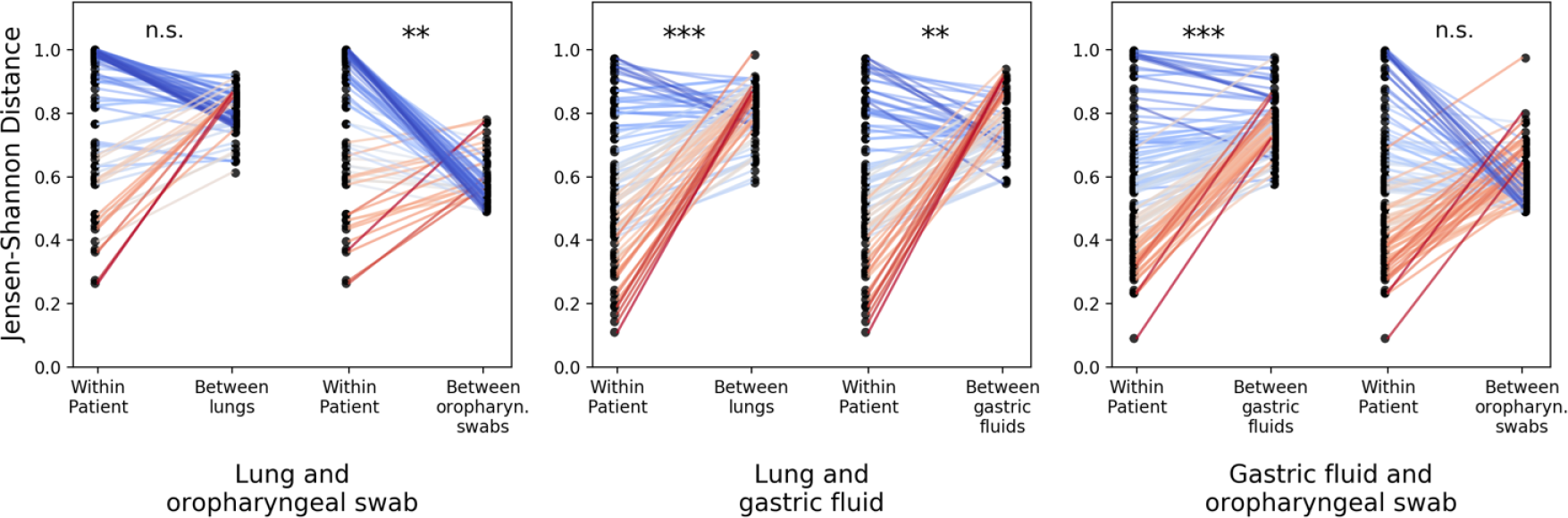
Lung and gastric microbial communities are driven primarily by person rather than body site. We compared the within-patient JSD for all pairs of aerodigestive sites with the average across-patient JSD between each of the sites in the within-patient comparison. Each panel shows different aerodigestive pairs; the two slope graphs correspond to different across-site comparisons; and each point corresponds to one patient. The left points in each slope graph show the within-patient JSD for the respective pair of sites for each patient (and are the same within each panel). The right points show the average JSD between the corresponding patient’s site X and all other patients’ site X. For example, the middle panel shows that the JSD between lung and gastric fluid communities within patients is lower than the average JSD between different lungs (left slope graph) and the average JSD between different gastric fluid samples (right slope graph). P values were calculated with a Wilcoxon signed-rank p-values using Python’s `scipy.stats.wilcoxon` function. * * * : *p* < 10^−10^; ** : 10^−10^ < *p* < 10^−7^, table of comparisons and p-values can be found in Supplementary Table 3.

### 4.3 Aspiration modulates the relationship between lung and oropharyngeal microbiomes but not the lung and stomach

Next, we investigated the impact of oropharyngeal dysphagia and aspiration on the aerodigestive microbiome. To assess whether there were large-scale differences in the microbiomes of aspirators and non-aspirators, we compared the alpha diversity for each aerodigestive site between these patient groups. Aspirators did not have significantly different alpha diversity in any of the aerodigestive sites for any of the metrics we compared (Supplementary Figure 3). Next, we attempted to identify individual OTUs which were differentially abundant between aspirators and non-aspirators. No OTUs or genera were significant in any aerodigestive site after correcting for multiple tests (Supplementary Table 4).

We next leveraged our within-patient sampling to investigate the effect of aspiration on the relationships between sites in the aerodigestive tract. Aspirators had significantly more similar lung and oropharyngeal communities than non-aspirators (Figure 5A, p = 0.04) and were much more likely to have the pre-defined oropharyngeal-lung microbes in both their oropharynx and lungs than non-aspirators (p = 4×10^−5^) (Figure 5B). Lung-oropharynx exchanged OTUs co-occurred in a median of 40% of aspirators’ lung and oropharyngeal communities but only 17% of non-aspirators’. Aspirators were not more likely to have stomach-lung microbes present in both the lungs and gastric fluid than non-aspirators (Figure 4B, p = 0.5), and lung and gastric communities of aspirating patients were not necessarily more similar to each other than those of non-aspirating patients (Figure 4A, p = 0.6).

**Figure 5:**
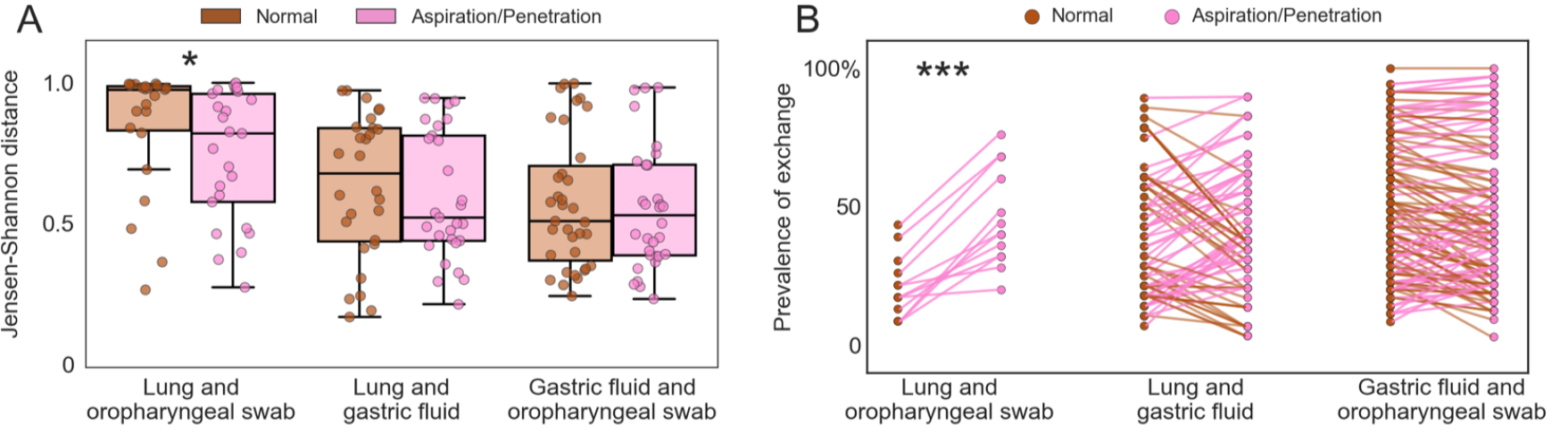
Dysphagia increases aspiration of microbes from the oropharynx but not the stomach. (A) Intra-patient Jensen Shannon distance for different aerodigestive site comparisons in non-aspirators (brown) and aspirators (pink). Each point represents one patient. P-values (Wilcoxon rank sums test, calculated with Python’s `scipy.stats.ranksums` function): lung and oropharyngeal swab *p* = 0.04, lung and gastric fluid *p* = 0.5, gastric fluid and oropharyngeal swab *p* = 0.8. (B) Percentage of patients with the previously defined exchanged microbes present in both of the respective sites (x-axis) in non-aspirators (brown) and aspirators (pink). Each pair of points represents one exchanged OTU. P-values (paired t-test on *log*10 prevalence values, calculated with Python’s `scipy.stats.ttest_rel` function: lung and oropharyngeal swab *p* = 4 × 10^−5^, lung and gastric fluid *p* = 0.5, gastric fluid and oropharyngeal swab *p* = 0.1.

To identify potential microbial biomarkers of aspiration, we looked at the exchanged OTUs which were most frequently present in the lung and oropharyngeal communities of aspirators relative to non-aspirators. In the oropharyngeal-lung exchanged OTUs, these were an unknown OTU in the *Flavobacteriaceae* family, OTUs in the *Fusobacterium*, *Rothia*, *Veillonella* genera, and an unknown OTU in the *Prevotellaceae* family, among others (Table 3, gastric-lung OTUs in Supplementary Table 5).

**Table 3:** Prevalence of lung-oropharynx exchanged OTUs. Prevalence is calculated as the percentage of patients who have the OTU present in both their lungs and oropharynx, calculated separately among aspirators (N = 25) and non-aspirators (N = 23). OTUs are ordered by their differential prevalence in aspirators relative to non-aspirators, and are labeled with their family- and genus-level taxonomies. Blank genus names indicate OTUs which were not annotated at the genus level. A similar table for the lung-gastric exchange OTUs can be found in Supplementary Table 1.

| Family | Genus | Non-aspirator | Aspirator | Difference |
| --- | --- | --- | --- | --- |
| Flavobacteriaceae |  | 8.7 | 48.0 | 39.3 |
| Fusobacteriaceae | Fusobacterium | 30.4 | 68.0 | 37.6 |
| Micrococcaceae | Rothia | 8.7 | 44.0 | 35.3 |
| Veillonellaceae | Veillonella | 26.1 | 60.0 | 33.9 |
| Prevotellaceae |  | 43.5 | 76.0 | 32.5 |
| Porphyromonadaceae | Porphyromonas | 39.1 | 68.0 | 28.9 |
| Streptococcaceae | Streptococcus | 13.0 | 40.0 | 27.0 |
| Veillonellaceae | Centipeda | 8.7 | 32.0 | 23.3 |
| Prevotellaceae | Prevotella | 17.4 | 36.0 | 18.6 |
| Leptotrichiaceae | Streptobacillus | 21.7 | 40.0 | 18.3 |
| Fusobacteriaceae | Fusobacterium | 17.4 | 32.0 | 14.6 |
| Aerococcaceae | Abiotrophia | 21.7 | 28.0 | 6.3 |
| Neisseriaceae | Neisseria | 17.4 | 20.0 | 2.6 |

We used Random Forest classifiers trained on the presence of exchanged OTUs in different sites and on the entire aerodigestive communities in order to test their potential as diagnostics for aspiration. We evaluated these classifiers by calculating the Fisher’s exact p-values and the area under the ROC curve (AUC) on leave-one-out predictions, where an AUC of 1.0 indicates a perfect classifier and an AUC of 0.5 is a classifier which assigns labels randomly [34]. The concordant presence or absence of exchanged OTUs in the two sites slightly improved classifiers based on the oropharyngeal-lung OTUs but not the ones based on the lung-gastric OTUs, relative to classifiers based on the presence of the exchanged OTUs in either site alone (Table 4; classifiers trained on the abundance of exchanged OTUs presented in Supplementary Table 6). However, these marginal results suggest that additional work will be necessary to develop these exchanged OTUs into reliable diagnostic biomarkers.

**Table 4:** Classifiers based on the presence of exchanged OTUs. (Top) Classifiers built from the presence of lung-oropharynx exchanged OTUs. (Bottom) Classifiers built from the presence of lung-gastric exchanged OTUs. Rows indicate which microbial community was used to train each classifier. In the “concordance” classifiers, OTUs which were either present or absent in both sites were coded as 1 and OTUs which were present in one site but absent in the other were coded as 0. AUCs are calculated as the area under the ROC curve from leave-one-out predictions. Fisher’s exact p values are calculated on the leave-one-out predictions. Similar classifiers built from the abundance of exchanged OTUs are shown in Supplementary Table 6.

| **Lung-oropharynx OTUs (13)** | AUC | p | N (non-asp/asp) |
| --- | --- | --- | --- |
| Lung | 0.66 | 0.08 | 33/33 |
| Oropharyngeal | 0.57 | 0.35 | 43/36 |
| Concordance | 0.63 | 0.05 | 23/25 |
| **Lung-gastric OTUs (76)** |  |  |  |
| Lung | 0.60 | 0.14 | 33/33 |
| Gastric fluid | 0.65 | 0.03 | 48/41 |
| Concordance | 0.53 | 1.0 | 28/29 |

**Table 5:** Classifiers based on perturbed relationship between lung and oropharyngeal microbiota can distinguish aspirators from non-aspirators. Areas under the ROC curve (AUC) and Fisher p-values calculated from classifiers trained on the entire microbial communities. Each row is a different classifier based on different combinations of aerodigestive communities, indicated in the “Sites” column. In the multi-site classifiers, the abundances of OTUs in different sites were used as separate features. AUCs and Fisher’s p values were calculated from the leave-one-out predictions for each sample.

| Sites | AUC | Fisher p-value | N (non-asp/asp) |
| --- | --- | --- | --- |
| Lung | 0.63 | 0.32 | 33/33 |
| Oropharyngeal swab | 0.69 | 0.02 | 43/36 |
| Gastric fluid | 0.66 | 0.08 | 48/41 |
| Lung and oropharyngeal swab | 0.79 | 0.01 | 23/25 |
| Lung and gastric fluid | 0.66 | 0.02 | 28/29 |
| Oropharyngeal swab and gastric fluid | 0.73 | 0.003 | 35/32 |
| All three sites | 0.78 | 0.01 | 19/23 |

**Table 6:** Reflux characteristics for 28 patients measured by pH-MII.

|  | Mean (std) |
| --- | --- |
| Number of acid episodes | 23.1 (25.5) |
| Number of nonacid episodes | 15.4 (16.4) |
| Number of pH only episodes | 16.3 (12.5) |
| Number of total reflux episodes | 38.0 (32.4) |
| Percent time proximal reflux | 0.48 (0.46) |
| Percent time distal reflux | 1.3 (1.1) |
| Percent time pH < 4 | 5.4 (5.2) |
| Number abnormal by pH-metry | 9/28 |
| Number abnormal by MII | 3/28 |

Using Random Forest classifiers trained on the entire microbiomes, we found that combining the oropharynx and lung communities resulted in a better classifier than either community alone (Table 6). Surprisingly, the classifiers trained on oropharyngeal and gastric communities performed well, despite our expectation that aspiration-induced changes in the microbiome would manifest in the lungs rather than the oropharynx or stomach. We confirmed that the patients’ aspiration status was not confounded with proton pump inhibitor usage (Fisher exact p-value = 0.8, Supplementary Table 1), but there may be other co-morbidities or unmeasured confounders that could be driving the differences detected in these communities. However, taken together, these results suggest that identifying a biomarker for aspiration based on bacteria in both the lungs and oropharynx may be possible, and that these two sites together contain more information about a patient’s aspiration status than either site alone.

### 4.4 Reflux may impact the relationship between lung and stomach microbiomes

Reflux profiles for the 28 patients are shown in Table 6. The percent of full column, distal, and proximal reflux were slightly negatively correlated with gastric-lung JSD, indicating that patients with more frequent reflux may have more similar gastric and lung microbial communities (Figure 6). However, the large range of gastric-lung JSDs across all patients and relatively weak correlation suggests that other non-reflux factors likely contribute more to the similarities between gastric and lung communities that are observed across all people. Similarly, we were not able to identify relationships between gastric-lung JSD and PPI usage (Supplementary Figures 9 and 10).

**Figure 6:**
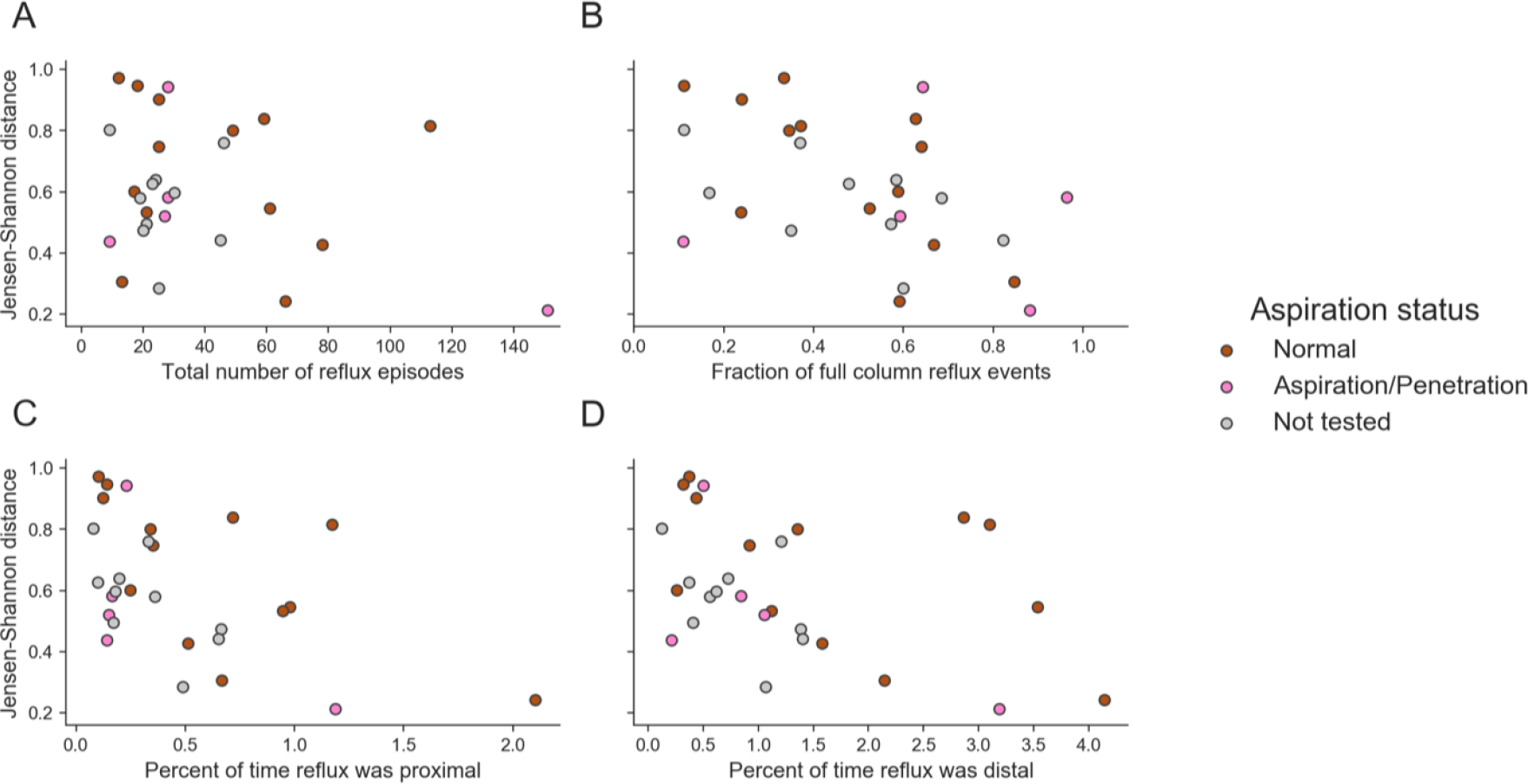
Reflux severity may correlate with the similarity between lung and gastric communities. Each plot shows the correlation between different reflux measures and the within-patient Jensen-Shannon distance between BAL and gastric fluid samples. Points are colored according to aspiration status. All reflux measures include both acid- and non-acid reflux. Spearman correlation and p-values: total number of reflux episodes *ρ*_*s*_ = −0.25, *p* = 0.2, percentage of full column reflux events *ρ*_*s*_ = −0.40, *p* = 0.04, percent of time reflux was proximal *ρ*_*s*_ = −0.55, *p* = 0.002, percent of time reflux was distal *ρ*_*s*_ = −0.45, *p* = 0.02.

## 5 Discussion

In this study, we characterized the relationships between the oropharyngeal, lung, and gastric microbiomes in a large pediatric cohort with and without swallowing dysfunction. Leveraging our simultaneous sampling of multiple sites per patient, we find that there exists a “core” oropharyngeal microbiome across patients, but lung and gastric communities vary and are distinct to individuals. Within patients, lung and oropharyngeal communities remain most distinct. We show for the first time that in patients with impaired swallowing, the lung microbiome shifts toward oropharyngeal flora rather than gastric flora. Our results also suggest that identifying biomarkers for aspiration based on the presence of certain bacteria in both the lungs and oropharynx may ultimately be possible.

There are several limitations to our study. First, because it is unethical to perform bronchoscopies on healthy children, our patients in this study had respiratory symptoms. Furthermore, these patients were on variety of medications (Table 1), which may affect microbial community compositions and relationships. However, we believe that our patient population represents patients typically seen in aerodigestive centers and that understanding the degree of microbial exchange is most clinically relevant in patients with symptoms. The microbial populations we found in this study are similar to those of previously published studies of both healthy and symptomatic adults which reinforces the validity of our results [12, 13, 17, 18]. We also confirmed that medication use and symptoms were not confounded with aspiration status (Supplementary Table 1). Second, the number of patients undergoing pH-MII testing was relatively small which limits our conclusions about the impact of gastroesophageal reflux on the lung. However, our study raises enough concerns about the significance of oropharyngeal-lung exchange in children with impaired swallowing that gastroesophageal reflux should not be considered as the primary source of microbial exchange causing pulmonary symptoms. Third, the diagnosis of oropharyngeal dysphagia in this study was based on VFSS. While this only categorizes patients based on a “one-point-in-time” study, it is the gold standard test to diagnose oropharyngeal dysphagia in children and therefore we feel it is appropriate for use in this study. Finally, the low biomass of BAL and gastric fluid samples could lead to sequencing artifacts or contamination. We did not explicitly remove potential background environmental or sampling sequences from our data, though our sampling methods was carefully developed in order to minimize potential contaminants [12, 16]. The low biomass of BAL and gastric fluid samples also resulted in fewer total sequencing reads than the oropharyngeal swabs (Supplementary Figure 11), perhaps contributing partially to the high variability we observed between these communities. However, many of our conclusions depend upon within-patient analyses, which reduce spurious results.

Despite these limitations, our findings have broad clinical implications for the understanding and treatment of oropharyngeal dysphagia with resultant aspiration. Our clinical finding that the lung microbiome in children with aspiration shifts toward the oropharynx rather than the stomach highlights the importance of understanding the primary driver of microbial exchange so that therapies can be tailored accordingly. For example, if the mechanism of lung symptoms and disease in aspirating children results from a microbial shift towards oropharyngeal flora, anti-reflux surgery will be of no benefit to preventing oropharyngeal-lung exchange. Instead, therapies may need to be tailored to focused on changing oropharyngeal flora or salivary properties.

While there are no existing pediatric microbiome studies of the aerodigestive microbiome in patients with dysphagia, there is evidence that children with oropharyngeal dysphagia are predisposed to pneumonia and that this could be due to increased aspiration of microbes from the oral microbiome. In a study of 382 children undergoing VFSS, evidence of aspiration predicted pneumonia risk, though the causative organisms for these pneumonias were not known [35]. In cohort of elderly aspirating patients, oral colonization by respiratory pathogens was associated with increased risk of pneumonia, highlighting the potential importance of oral flora in influencing the lung outcomes [36]. Finally, a previous study of healthy adults found that individuals with oropharyngeal bacteria in their lungs had increased evidence of inflammatory metabolomic signals, suggesting that even a change of lung flora to commensal oropharyngeal bacteria can trigger inflammation even in healthy patients [22]. Our results add to these findings and suggest that changes in the lung microbiome towards oropharyngeal flora merit additional study to determine if these shifts result in increased morbidity or worse clinical outcomes, including the development of pneumonia.

From a microbial perspective, we identified bacterial families and genera that are more commonly exchanged between the oropharynx and lungs of children that aspirate than of children with intact swallowing mechanism. While there are no other 16S sequencing studies determining aspiration pneumonia risk in children, there is evidence from the adult literature that similar bacteria are involved in aspiration pneumonia risk. For example, oropharyngeal *Streptococci* were found to be more abundant in the lungs of adults with pneumonia and aspiration risk factors than without aspiration risk [37]. In a study of 173 adults in long term care facilities, patients with oropharyngeal *Prevotella* and *Veillonella* had increased risk of death from pneumonia compared to patients who had oropharyngeal *Neisseria* and *Fusobacterium* [38]. Our study is a critical first step toward identifying bacteria present in the oropharynx and lungs of aspirating children that may result in higher risk for pneumonias, with additional studies needed to determine their impact on pediatric outcomes.

In summary, our findings suggest that interventions to reduce aspiration-related respiratory complications due to increased microbial exchange should target aspiration from the oropharynx rather than the stomach. This microbial data supports the clinical observation that antireflux surgery fails to prevents pulmonary complications such as pneumonias or hospitalizations [3, 7, 8, 9, 10, 11]. By simultaneously sampling multiple sites per patient, we show that the lung and stomach microbiomes are highly variable across patients and determined primarily by patient rather than body site. If aerodigestive microbial communities are indeed specific to each individual, interventions targeting the aerodigestive microbiome may benefit from personalized medicine approaches. Finally, understanding the relationships between aerodigestive communities in aspirating and non-aspirating patients provides insight into the potential pathophysiology behind aspiration-related respiratory outcomes and suggests potential diagnostics and therapeutics for future investigation.

## 6 Declarations

### 6.1 Acknowledgments

We thank Scott Olesen and Manu Kumar for helpful discussions on statistical analyses, and Nathaniel Chu and members of the Alm lab for helpful discussions on presenting and visualizing the results.

### 6.3 Author contributions

R.R. designed the study, recruited patients, and performed the endoscopies. R.R. led patient recruitment. A.L. and S.I. assisted with patient recruitment. K.L. performed and interpreted the videofluoroscopic swallow studies. K.M. performed the bronchoscopies in this study. K.F. and S.S. performed the DNA isolation for 16S sequencing. C.D. processed and analyzed the microbiome data. C.D., E.A., and R.R. interpreted the results. C.D. and R.R. wrote the manuscript.

### 6.4 List of abbreviations

EGD: esophagogastroduodenoscopy
BAL: bronchoalveolar lavage
MII: multichannel intraluminal impedance
VFSS: videofluoroscopic swallow study
OTU: operational taxonomic unit
JSD: Jensen-Shannon distance
PERMANOVA: permutational multivariate analysis of variance
AUC: area under the ROC (receiver operating characteristic) curve

