## Supplementary Tables and Figures for "Aerodigestive sampling reveals altered microbial exchange between lung, oropharyngeal, and gastric microbiomes in children with impaired swallow function"

### 1 Supplementary Tables and Figures

#### 1.1 Supplementary Tables

| | Aspiration/Penetration | Normal | Fisher's p |
|---|---|---|---|
| **Demographics** | | | |
| Gender | 30 M, 17 F | 31 M, 26 F | |
| Age | 3.9 ± 4.1 years | 6.7 ± 4.7 years | |
| **Symptom and quality of life scores** | | | |
| PGSQ symptom score | 0.9 ± 0.75 | 0.9 ± 0.71 | |
| PGSQ total score | 0.7 ± 0.61 | 0.9 ± 0.70 | |
| **Medications** | | | |
| Currently taking PPIs | 55% (26/47) | 58% (33/57) | 0.84 |
| Currently taking H2 blockers | 15% (7/47) | 21% (12/57) | 0.46 |
| Current use of inhaled steroids | 72% (34/47) | 75% (43/57) | 0.82 |
| **Symptoms within last 6 months** | | | |
| Problem swallowing | 7% (3/41) | 22% (12/54) | 0.09 |
| Food stuck | 29% (12/41) | 26% (14/54) | 0.82 |
| Difficulty swallowing | 34% (14/41) | 28% (15/54) | 0.51 |
| Abdominal pain | 37% (16/43) | 32% (18/56) | 0.67 |
| Constipation | 35% (14/40) | 37% (19/52) | 1.00 |
| Weight loss | 22% (9/40) | 30% (16/54) | 0.49 |
| Food coming up | 30% (13/44) | 39% (22/56) | 0.40 |
| Chest pain | 10% (4/40) | 24% (13/54) | 0.11 |
| Chronic cough | 65% (30/46) | 72% (38/53) | 0.52 |
| **Infection history within 6 months** | | | |
| History of pneumonia | 33% (15/45) | 33% (18/54) | 1.00 |
| Recent history of ear infection | 42% (18/43) | 19% (10/54) | 0.01 |
| Recent history of sinus infection | 25% (11/44) | 26% (14/53) | 1.00 |
| History of any recent antibiotics | 28% (13/47) | 39% (22/57) | 0.30 |

Supplementary Table 1: Patient demographics, separated by aspiration status. While all patients were given questionnaires, not all patients completed the answers to all questions. Fisher's exact p-values were calculated on the contingency table of aspiration status and the metadata indicated in each row, and tests whether the distribution of symptoms, medications, or infection history is similarly distributed between aspirators and non-aspirators. For example, the Fisher's exact test for the "Currently on PPIs" column was calculated on the following contingency table: [[(47 - 26), 26], [(57 - 33), 33]].

| | Aspiration/penetration | Normal | Not tested |
|---|---|---|---|
| BAL, gastric fluid, and oropharyngeal swab | 23 | 19 | 24 |
| BAL and gastric fluid | 6 | 9 | 7 |
| BAL and oropharyngeal swab | 2 | 4 | 1 |
| Gastric fluid and oropharyngeal swab | 9 | 16 | 20 |
| Oropharyngeal swab and stool | | | 20 |
| BAL only | 2 | 1 | 3 |
| Gastric fluid only | 3 | 4 | 5 |
| Oropharyngeal swab only | 2 | 4 | 31 |
| Stool only | | | 5 |
| Total patients | 47 | 57 | 118 |

Supplementary Table 2: Number of patients with each combination of body sites sequenced, separated by aspiration status.

| Within-patient sites | Between-patient site | p | Direction |
|---|---|---|---|
| Lung and oropharynx | lung | 0.71 | within < between |
| Lung and oropharynx | oropharynx | $2 \times 10^{-8}$ | within > between |
| Lung and gastric fluid | lung | $9 \times 10^{-12}$ | within < between |
| Lung and gastric fluid | gastric fluid | $7 \times 10^{-8}$ | within < between |
| Gastric fluid and oropharynx | gastric fluid | $5 \times 10^{-15}$ | within < between |
| Gastric fluid and oropharynx | oropharynx | 0.18 | within < between |

Supplementary Table 3: **Lung and gastric microbial communities are driven primarily by person rather than body site.** We compared the within-patient JSD for all pairs of aerodigestive sites with the average across-patient JSD between each of the sites in the within-patient comparison. For each pair of aerodigestive sites, we compared each patient's within-patient JSD with the average across-patient JSD for each site in the pair. For example, the top row shows the comparisons between (1) the JSD between each patient's own oropharyngeal and lung communities ("Within-patient sites") with (2) the average JSD between that patient's lung community and all other lung communities ("Between-patient site"). We calculated Wilcoxon signed-rank p-values using Python's `scipy.stats.wilcoxon` function ("p"). The "Direction" column indicates whether the within-patient JSD was larger (i.e. more different) than the between-patient JSD (`within > between`), or vice-versa (`within < between`).

| | Site | Minimum q-value |
|---|---|---|
| **Genus-level** | Lung | 0.23 |
| | Gastric fluid | 0.35 |
| | Oropharyngeal swab | 0.27 |
| **OTU-level** | Lung | 0.60 |
| | Gastric fluid | 0.11 |
| | Oropharyngeal swab | 0.18 |

Supplementary Table 4: Differential abundance analysis comparing aspirators vs. non-aspirators yields no significant results. P-values were calculated on the relative abundances of genera and OTUs with the Kruskal-Wallis test implemented in Python's `scipy.stats.mstats` module (function `kruskalwallis`, a non-parametric test and an implementation which accounts for ties). P-values were corrected for multiple hypothesis testing with the `multipletests` function from `statsmodels.sandbox.stats.multicomp`, with the Benjamini/Hochberg correction (`method = 'fdr_bh'`). P-values were corrected separately for each site and each level of analysis (e.g. OTU or genus-level).

| Family | Genus | Non-aspirator | Aspirator | Difference |
|---|---|---|---|---|
| Neisseriaceae | Neisseria | 7.1 | 41.4 | 34.2 |
| Porphyromonadaceae | Porphyromonas | 28.6 | 62.1 | 33.5 |
| Pasteurellaceae | Haemophilus | 50.0 | 82.8 | 32.8 |
| Lachnospiraceae | Coprococcus | 10.7 | 37.9 | 27.2 |
| Micrococcaceae | Rothia | 14.3 | 41.4 | 27.1 |
| Prevotellaceae | Prevotella | 25.0 | 51.7 | 26.7 |
| Carnobacteriaceae | Granulicatella | 32.1 | 58.6 | 26.5 |
| Bacillales_Incertae_Sedis_XI | Gemella | 42.9 | 69.0 | 26.1 |
| Pasteurellaceae | Haemophilus | 57.1 | 82.8 | 25.6 |
| Actinomycetaceae | Actinomyces | 17.9 | 41.4 | 23.5 |
| Streptococcaceae | Streptococcus | 39.3 | 62.1 | 22.8 |
| Lachnospiraceae | Oribacterium | 14.3 | 34.5 | 20.2 |
| Leptotrichiaceae | Streptobacillus | 17.9 | 37.9 | 20.1 |
| Lachnospiraceae | Lachnoanaerobaculum | 17.9 | 37.9 | 20.1 |
| Fusobacteriaceae | Fusobacterium | 42.9 | 62.1 | 19.2 |
| Prevotellaceae | | 50.0 | 69.0 | 19.0 |
| Flavobacteriaceae | Planobacterium | 14.3 | 31.0 | 16.7 |
| Leptotrichiaceae | Leptotrichia | 14.3 | 31.0 | 16.7 |
| Erysipelotrichaceae | Solobacterium | 17.9 | 34.5 | 16.6 |
| Prevotellaceae | Prevotella | 21.4 | 37.9 | 16.5 |
| Pasteurellaceae | Haemophilus | 28.6 | 44.8 | 16.3 |
| Veillonellaceae | Veillonella | 35.7 | 51.7 | 16.0 |
| Enterobacteriaceae | Escherichia/Shigella | 46.4 | 62.1 | 15.6 |
| Prevotellaceae | | 46.4 | 62.1 | 15.6 |
| Neisseriaceae | Neisseria | 60.7 | 75.9 | 15.1 |
| Streptococcaceae | Streptococcus | 75.0 | 89.7 | 14.7 |
| Veillonellaceae | Veillonella | 35.7 | 48.3 | 12.6 |
| Prevotellaceae | Prevotella | 42.9 | 55.2 | 12.3 |
| Micrococcaceae | Rothia | 42.9 | 55.2 | 12.3 |
| Streptococcaceae | Streptococcus | 42.9 | 55.2 | 12.3 |
| Prevotellaceae | Prevotella | 64.3 | 75.9 | 11.6 |
| Unknown Burkholderiales | | 10.7 | 20.7 | 10.0 |
| Bacteroidaceae | Bacteroides | 14.3 | 24.1 | 9.9 |
| Porphyromonadaceae | Porphyromonas | 21.4 | 31.0 | 9.6 |
| Moraxellaceae | Moraxella | 39.3 | 48.3 | 9.0 |
| Prevotellaceae | Prevotella | 57.1 | 65.5 | 8.4 |
| Leptotrichiaceae | Leptotrichia | 21.4 | 27.6 | 6.2 |
| Fusobacteriaceae | Fusobacterium | 25.0 | 31.0 | 6.0 |
| Porphyromonadaceae | Porphyromonas | 50.0 | 55.2 | 5.2 |
| Neisseriaceae | Neisseria | 17.9 | 20.7 | 2.8 |
| Veillonellaceae | Veillonella | 89.3 | 89.7 | 0.4 |
| Coriobacteriaceae | Atopobium | 21.4 | 20.7 | -0.7 |
| Unknown Bacteria | | 21.4 | 20.7 | -0.7 |
| Enterococcaceae | | 85.7 | 82.8 | -3.0 |
| Chloroplast | Streptophyta | 10.7 | 6.9 | -3.8 |
| Pasteurellaceae | Haemophilus | 17.9 | 13.8 | -4.1 |
| Unknown Bacillales | | 17.9 | 13.8 | -4.1 |
| Unknown Bacillales | | 17.9 | 13.8 | -4.1 |
| Lactobacillaceae | Lactobacillus | 28.6 | 17.2 | -11.3 |
| Pasteurellaceae | Haemophilus | 32.1 | 20.7 | -11.5 |
| Staphylococcaceae | Staphylococcus | 60.7 | 48.3 | -12.4 |
| Bacteroidaceae | Bacteroides | 17.9 | 3.4 | -14.4 |
| Comamonadaceae | Acidovorax | 17.9 | 3.4 | -14.4 |
| Porphyromonadaceae | Parabacteroides | 21.4 | 6.9 | -14.5 |
| Comamonadaceae | Pelomonas | 21.4 | 6.9 | -14.5 |
| Flavobacteriaceae | Chryseobacterium | 28.6 | 13.8 | -14.8 |
| Erysipelotrichaceae | Clostridium_XVIII | 21.4 | 3.4 | -18.0 |
| Lachnospiraceae | Ruminococcus2 | 25.0 | 6.9 | -18.1 |
| Flavobacteriaceae | Chryseobacterium | 50.0 | 31.0 | -19.0 |
| Neisseriaceae | Microvirgula | 57.1 | 37.9 | -19.2 |
| Enterobacteriaceae | Enterobacter | 82.1 | 62.1 | -20.1 |
| Mycobacteriaceae | Mycobacterium | 28.6 | 6.9 | -21.7 |
| Moraxellaceae | Acinetobacter | 60.7 | 37.9 | -22.8 |
| Streptococcaceae | Streptococcus | 60.7 | 37.9 | -22.8 |
| Bacteroidaceae | Bacteroides | 53.6 | 27.6 | -26.0 |
| Unknown Bacillales | | 57.1 | 31.0 | -26.1 |
| Moraxellaceae | Acinetobacter | 60.7 | 34.5 | -26.2 |
| Moraxellaceae | Enhydrobacter | 60.7 | 34.5 | -26.2 |
| Lactobacillaceae | Lactobacillus | 50.0 | 20.7 | -29.3 |
| Aeromonadaceae | Aeromonas | 57.1 | 27.6 | -29.6 |
| Moraxellaceae | Acinetobacter | 78.6 | 41.4 | -37.2 |
| Leuconostocaceae | Weissella | 78.6 | 37.9 | -40.6 |
| Moraxellaceae | Acinetobacter | 78.6 | 37.9 | -40.6 |
| Leuconostocaceae | Leuconostoc | 78.6 | 37.9 | -40.6 |
| Streptococcaceae | Lactococcus | 78.6 | 37.9 | -40.6 |
| Streptococcaceae | Lactococcus | 78.6 | 37.9 | -40.6 |

Supplementary Table 5: Prevalence of lung-gastric fluid exchanged OTUs. Prevalence is calculated as the percentage of patients who have the OTU present in both their lungs and oropharynx, calculated separately among aspirators (N = 29) and non-aspirators (N = 28). OTUs are ordered by their differential prevalence in aspirators relative to non-aspirators, and are labeled with their family- and genus-level taxonomies. Blank genus names indicate OTUs which were not annotated at the genus level.

| **Lung-oropharynx OTUs (13)** | AUC | p | N (non-asp/asp) |
|---|---|---|---|
| Lung | 0.59 | 0.21 | 33/33 |
| Oropharyngeal | 0.65 | 0.11 | 43/36 |
| Both | 0.70 | 0.08 | 23/25 |
| **Lung-gastric OTUs (76)** | | | |
| Lung | 0.56 | 0.45 | 33/33 |
| Gastric fluid | 0.66 | 0.03 | 48/41 |
| Both | 0.69 | 0.008 | 28/29 |

Supplementary Table 6: **Classifiers based on the abundance of exchanged OTUs.** (Top) Classifiers built from the abundance of lung-oropharynx exchanged OTUs. (Bottom) Classifiers built from the abundance of lung-gastric exchanged OTUs. Rows indicate which microbial community was used to train each classifier. In classifiers using two sites ("Both"), abundances of each exchanged OTU in each site were considered as separate features. AUCs are calculated as the area under the average ROC curve from leave-one-out predictions. Fisher's exact p values are calculated on the leave-one-out predictions using Python's `scipy.stats.fisher_exact` function.

### 1.2 Supplementary Figures

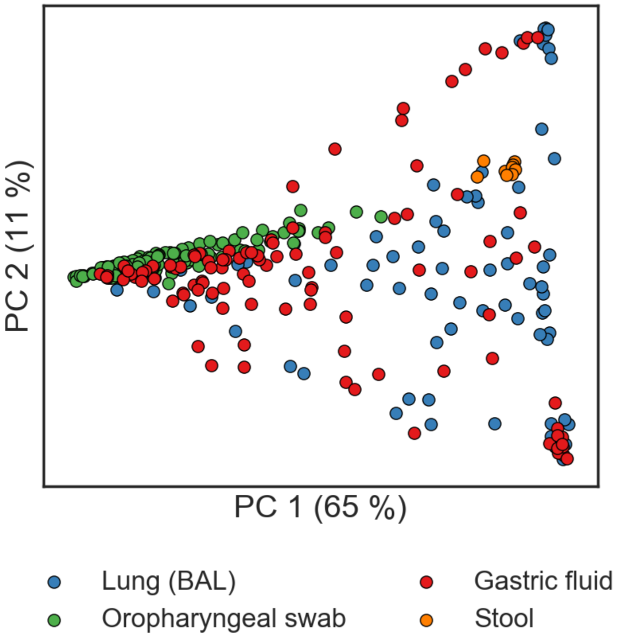


Supplementary Figure 1: PCoA plot of aerodigestive and stool microbial communities for patients in the sequencing batch not shown in the main text (N = 81 BAL, 124 oropharyngeal swab, 104 gastric fluid, and 11 stool samples). PERMANOVA on the BAL, gastric fluid, and oropharyngeal swab samples, $p = 0.001$.

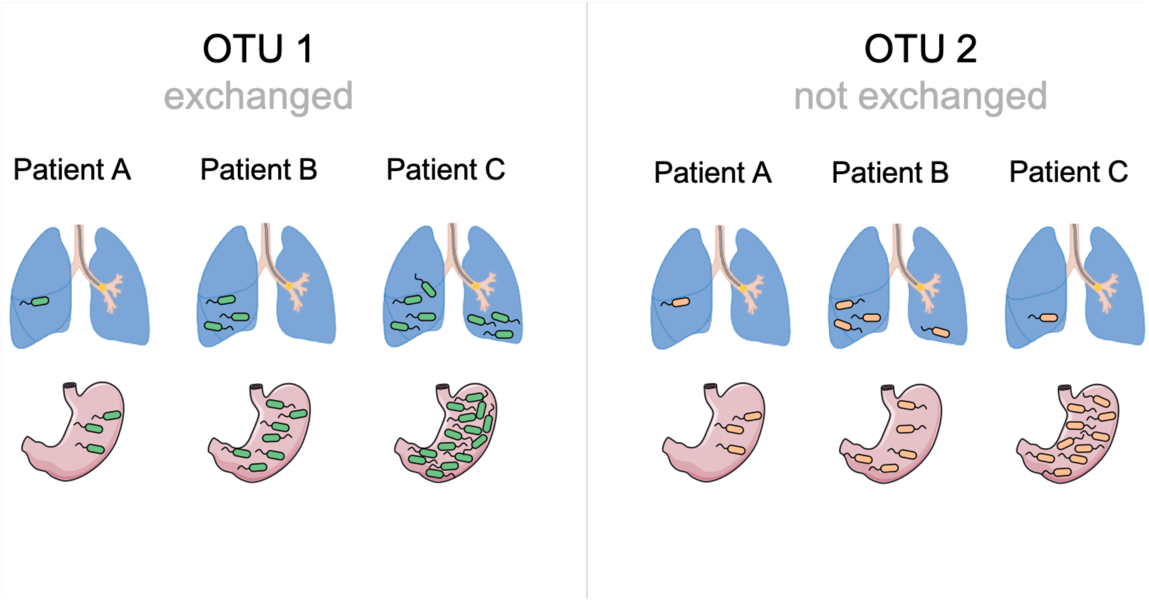


Supplementary Figure 2: Schematic illustrating an OTU which is considered exchanged between the lung and stomach (left) and one which is not (right). If an OTU is exchanged in two sites, its abundance in the two sites should be correlated across patients. For example, OTU 1 is exchanged across the lung and stomach. If its abundance in the stomach of Patient C is higher than in the stomach of Patient B, we expect that its abundance in the lungs of Patient C will be higher than in the lungs of Patient B. In contrast, OTU 2 is not exchanged, so knowing its abundance in Patient C's stomach relative to Patient B does not provide information about OTU 2's expected abundance in the lungs of Patient C. Lung image was adapted from Cancer Research UK / Wikimedia Commons and the stomach image is from Servier Medical Art.

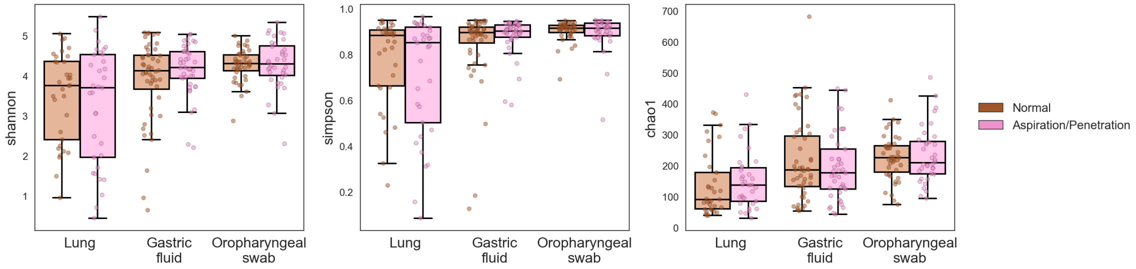


Supplementary Figure 3: Alpha diversity of each aerodigestive site compared between aspirators and non-aspirators. Each panel is a different metric, calculated with the respective metric (labeled on the y-axis) in `skbio.diversity.alpha`. All p-values for aspirator vs. non-aspirator comparisons are greater than 0.1 (Wilcoxon rank sums test calculated with Python's `scipy.stats.ranksums` function).

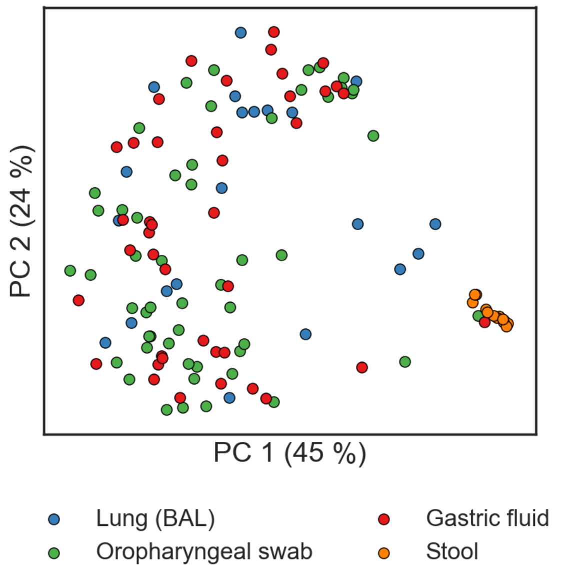


Supplementary Figure 4: PCoA plots of aerodigestive and stool microbial communities for all patients in the sequencing batch shown in Figure 2A, based on the Bray-Curtis distance.

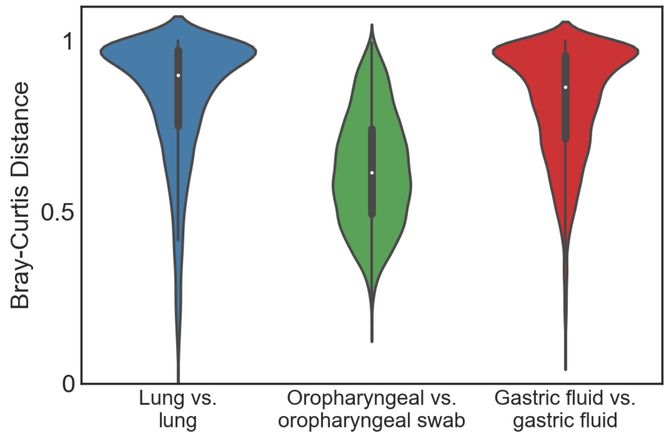


Supplementary Figure 5: Violin plots of the Bray-Curtis distance between samples from the same site across different patients.

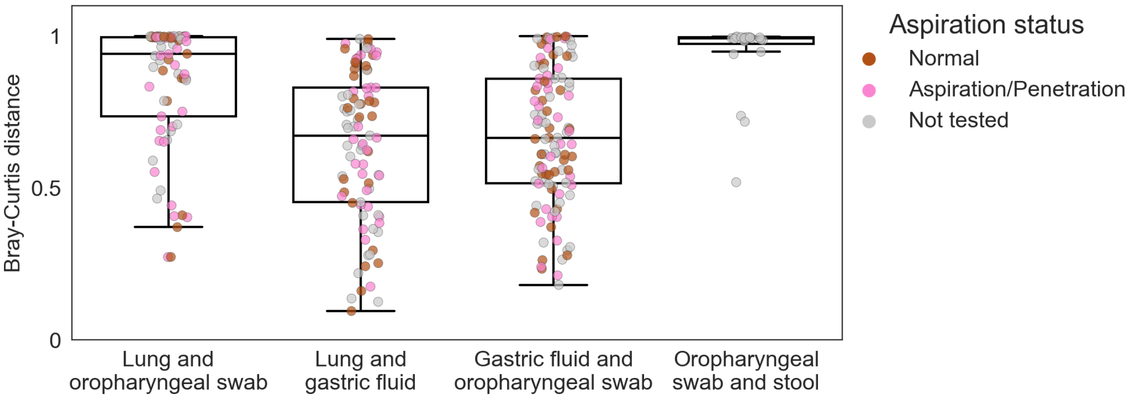


Supplementary Figure 6: Bray-Curtis distances between samples from different sites from the same patient. Comparisons between stool and oropharynx are included to contextualize these results, as these are expected to be very different. All comparisons are significant (Wilcoxon rank sums test calculated with Python's `scipy.stats.ranksums` function) except the lung and gastric fluid vs. gastric fluid and oropharyngeal swab beta diversities (p = 0.5) and the lung and oropharyngeal vs. oropharyngeal and stool (p = 0.2). All other comparisons: $p < 1 \times 10^{-6}$.

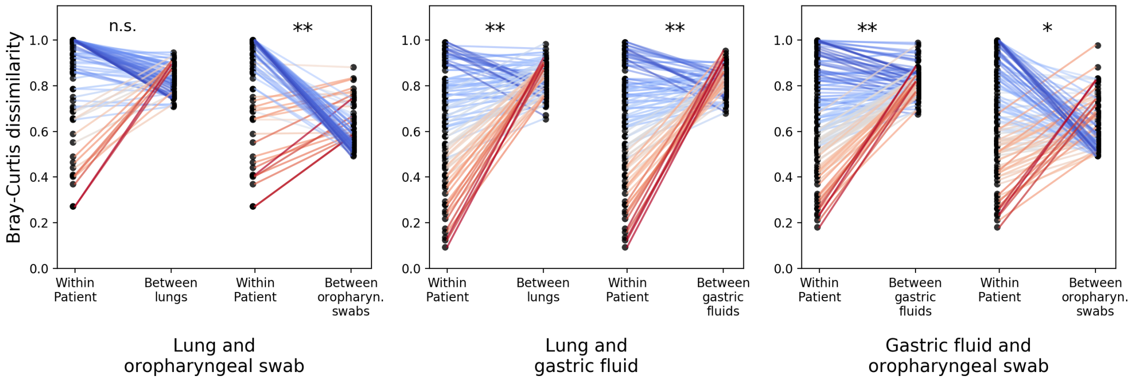


Supplementary Figure 7: Comparison between within-patient and between-patient beta diversities, as in Figure 4), calculated with the Bray-Curtis dissimilarity. $** : 10^{-10} < p < 10^{-6}$; $* : p = 0.04$.

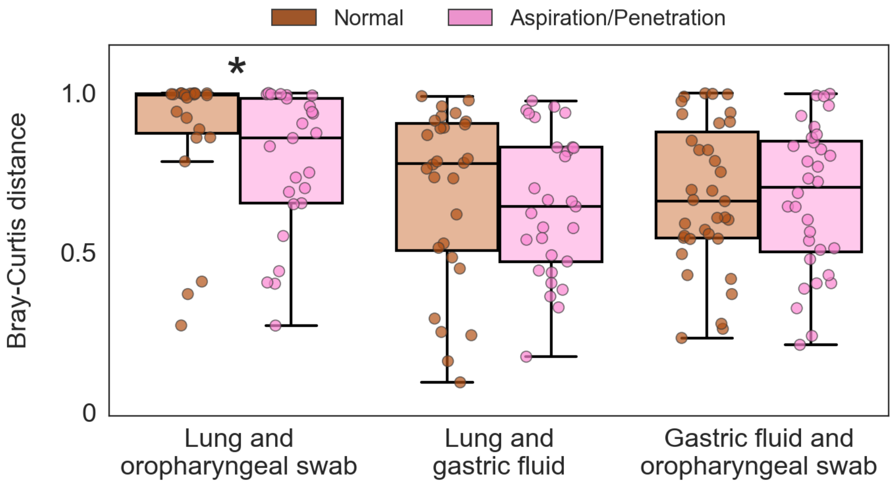


Supplementary Figure 8: Intra-patient Bray Curtis distance for different aerodigestive site comparisons in non-aspirators (brown) and aspirators (pink). Each point represents one patient. P-values (Wilcoxon rank sums test, calculated with Python's `scipy.stats.ranksums` function): lung and oropharyngeal swab $p = 0.02$, lung and gastric fluid $p = 0.5$, gastric fluid and oropharyngeal swab $p = 0.9$.

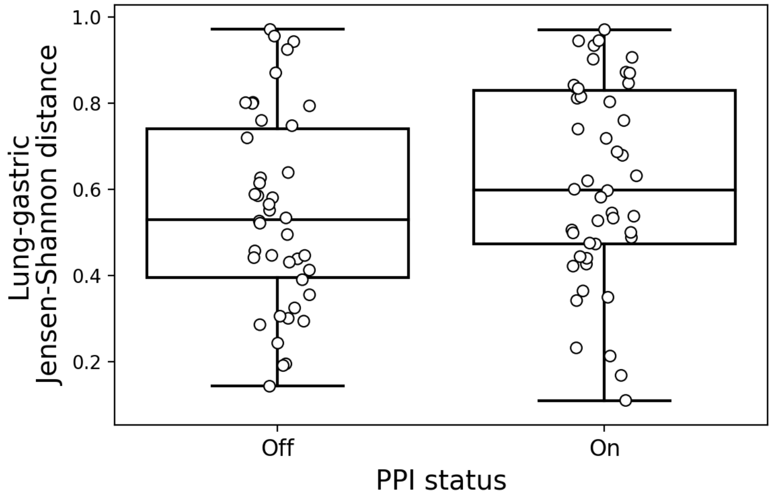


Supplementary Figure 9: Lung-gastric JSD vs. PPI status. Off PPI, N = 42; On PPI, N = 41. Wilcoxon rank sums test, calculated with Python's `scipy.stats.ranksums` function, p = 0.14.

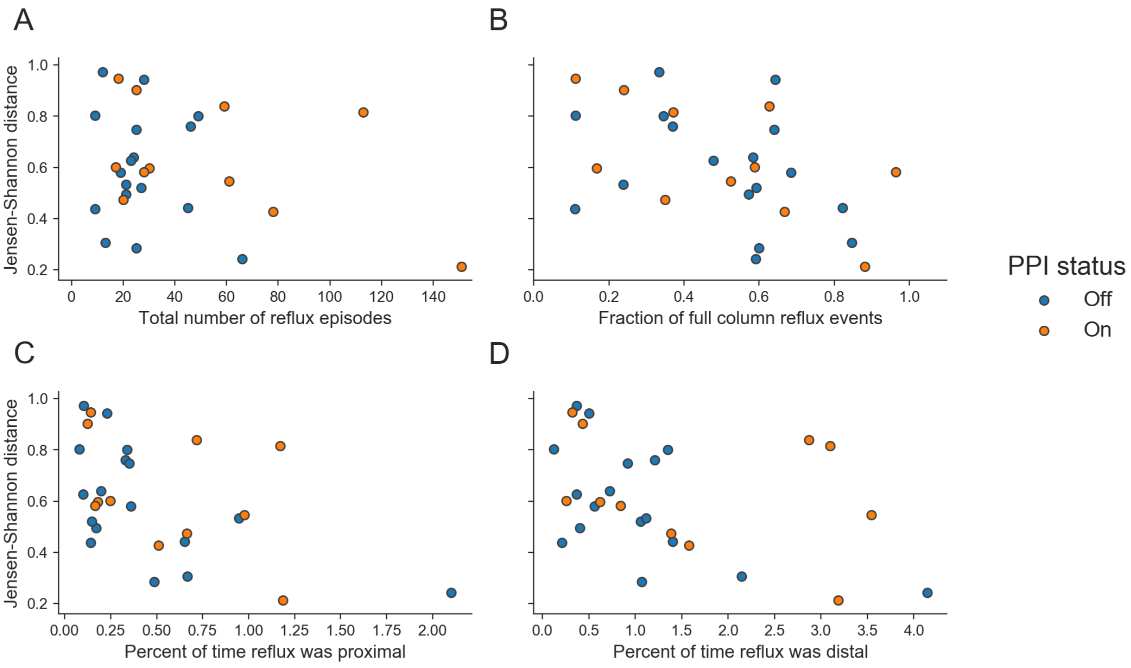


Supplementary Figure 10: Same data as in Figure 6, but points are colored by PPI status.

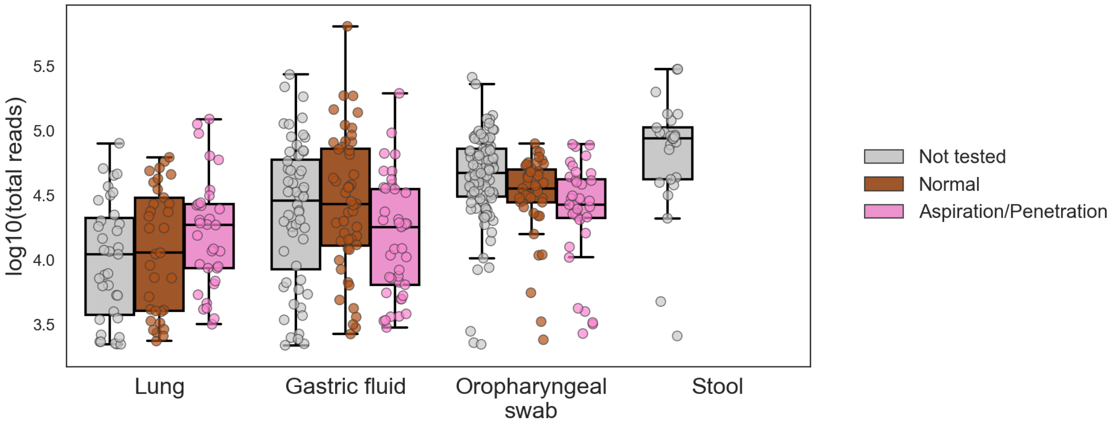


Supplementary Figure 11: Total reads per sample. P-values (Wilcoxon rank sums test, calculated with Python's `scipy.stats.ranksums` function) for aspiraton vs. non-aspirator comparison: lung $p = 0.3$, gastric fluid $p = 0.02$, oropharyngeal swab $p = 0.08$.
